# An active-matrix digital microfluidic platform for simultaneous short- and long-read viral genomic surveillance

**DOI:** 10.64898/2026.07.21.739725

**Authors:** Bingbing Zhang, Zhixin Fang, Qiming Liu, Jiajian Ji, Siyi Hu, Maolin Zhang, Yuebing Wang, Yong Chang, Xuhui Lai, Yanye Feng, Jiahao Li, Jun Yu, Chen Jiang, Arokia Nathan, Jinhua Li, Chao Yu, Hanbin Ma

**Author notes:** corresponding author. (Hanbin Ma); (Chao Yu); (Jinhua Li). these authors contributed equally to this work (Bingbing Zhang, Zhixin Fang).

## Abstract

The outbreak frequency and geographic distribution of viral pathogens are continuously expanding, making enhanced genomic surveillance an urgent global public health need. Parallel library preparation combining next-generation sequencing (NGS) and third-generation sequencing (TGS) can substantially improve the coverage and resolution of genomic surveillance, representing a key strategy for strengthening surveillance. Here we developed a complete sample-to-result system integrating a programmable active-matrix digital microfluidic (AM-DMF) chip with a bioinformatics analysis pipeline. Compared with conventional manual protocols used in public health laboratories, our system reduces reagent consumption by 72%, shortens library preparation time by 45% and decreases the inter-batch coefficient of variation (CV) by 20%. In 20 RT-qPCR-confirmed clinical samples, the system achieved complete concordance for viral identification and assigned serotypes/genotypes consistent with sequencing-based phylogenetic analysis. This system is field-deployable and enables rapid virus serotyping as well as in-depth genomic surveillance.

**Teaser:** A digital microfluidic platform integrating short- and long-read sequencing enables rapid comprehensive viral genome analysis.

## INTRODUCTION

The global burden of viral pathogens is escalating at an unprecedented pace. Beyond the recent COVID-19 pandemic, endemic and emerging viruses—including dengue, chikungunya, Zika, West Nile, influenza, Ebola, and Lassa—continue to expand their geographic ranges and clinical severity, driven by climate change, urbanization, and globalized travel(*1–3*). In this context, pathogen genomic surveillance has transitioned from a specialized research tool to an urgent public health imperative(*4–6*). Yet the most critical evolutionary events that shape outbreaks remain virtually invisible to conventional diagnostics. Resolving these highly dynamic processes requires simultaneously capturing the population-frequency distribution of viral variants and their single-molecule genomic architecture, a dual requirement that eludes current analytical paradigms.

NGS delivers the extreme fidelity and depth required for precise lineage assignment and low-frequency variant detection(*7*). However, its inherently short reads cannot determine whether distant mutations are physically linked on the same RNA molecule, nor can they reliably span highly structured or repetitive genomic segments, such as viral 5’ and 3’ untranslated regions(*8, 9*). TGS produces long reads capable of capturing near-full-length genomes, revealing large structural variations, and resolving haplotype phase, yet its elevated per-base error rate and quantitative biases render it insufficient as a standalone confirmatory tool(*10–12*). When deployed asynchronously, these two modalities offer incomplete, and often conflicting, portraits of the same infection. Strictly synchronous co-sequencing and co-analysis—leveraging TGS to map the macro-genomic structural backbone and NGS to rigorously error-correct and quantify micro-variant frequencies—are therefore essential to resolve complex evolutionary dynamics that neither platform can address alone(*13, 14*). Specifically, this unified approach provides the orthogonal validation required to determine whether distant resistance and immune-escape mutations are physically linked on the same virion, to distinguish genuine inter-serotype recombination from PCR-induced chimeric artifacts, and to accurately reconstruct the true haplotype architecture of minority variants destined to seed therapeutic failure(*15*).

Despite this scientific necessity, the current wet-laboratory and computational ecosystems remain fundamentally fragmented(*16*). RT-qPCR, the workhorse of clinical diagnostics, confirms infection within hours but yields no actionable intelligence on serotype, genotype, co-infection status, or critical mutations(*17*). NGS workflows necessitate centralized infrastructure, multi-hour library preparation, and substantial reagent volumes, rendering them incompatible with rapid frontline response. TGS can operate near the point of care but demands orthogonal validation before its outputs can drive public health action. Consequently, laboratories are forced to physically split precious, low-titer clinical specimens into entirely separate, temporally staggered workflows. This asynchronous segregation not only introduces temporal degradation bias and risks the severe stochastic dropout of rare variants, but ultimately destroys the spatiotemporal linkage of the highly unstable viral quasi-species. Commercial liquid-handling workstations merely mechanize these flawed serial protocols; they remain bulky, expensive, and unable to eliminate dead volumes or pipetting-path collisions during space-constrained multiplexing(*18*). While existing microfluidic devices have miniaturized library preparation, they operate exclusively on purified nucleic acids, as their microscale chambers cannot accommodate the macroscopic input volumes required for raw-sample extraction without sacrificing analytical sensitivity(*19–21*). Thus, few platforms can execute parallel, zero-cross-talk library preparation for both modalities starting directly from an unprocessed clinical aliquot. The bioinformatics bottleneck further exacerbates this diagnostic gap. True hybrid co-analysis of NGS and TGS data demands recursive reference anchoring, cross-platform error correction, haplotype reconstruction, and phylogenetic integration—workflows that conventionally require more than fifteen discrete command-line interventions and 30–60 minutes of specialist execution per sample. Furthermore, when fed with asynchronously prepared libraries, these algorithms are highly vulnerable to batch-effect artifacts, often misinterpreting sequencing noise as false-positive recombination events. Such complexity and unreliability are entirely incompatible with outbreak settings, where bioinformatic expertise is scarce and critical containment decisions must be rendered within hours(*22*).

Here, we report an integrated, sample-to-result architecture that overcomes these cross-cutting limitations (Fig.1). At its core is a programmable AM-DMF platform whose ultra-high-density electrode arrays enable software-defined, nanoliter-scale droplet manipulation across a macroscopic reaction area(*23–29*). By integrating an external reservoir that bridges the physical gap between large-volume sample input and microscale reaction capacity, the device executes fully automated nucleic acid extraction. Crucially, the platform then bifurcates the eluate into completely isolated reaction zones, achieving strictly synchronous parallel NGS and TGS library preparation from the same raw clinical aliquot without chemical cross-talk. By locking the dynamic viral cloud into a unified temporal snapshot, this physical architecture provides an isogenic ground truth for downstream analysis. The web-based autonomous pipeline leverages this synchronized input to execute deterministic multi-modal fusion—automatically performing intelligent recursive reference scanning, dual-platform alignment, zero-tolerance cross-validation of variants, consensus generation, and phylogenetic tracing. This framework compresses the entire hybrid co-analysis workflow from over 30 minutes to approximately 10 minutes.

**Fig. 1.**
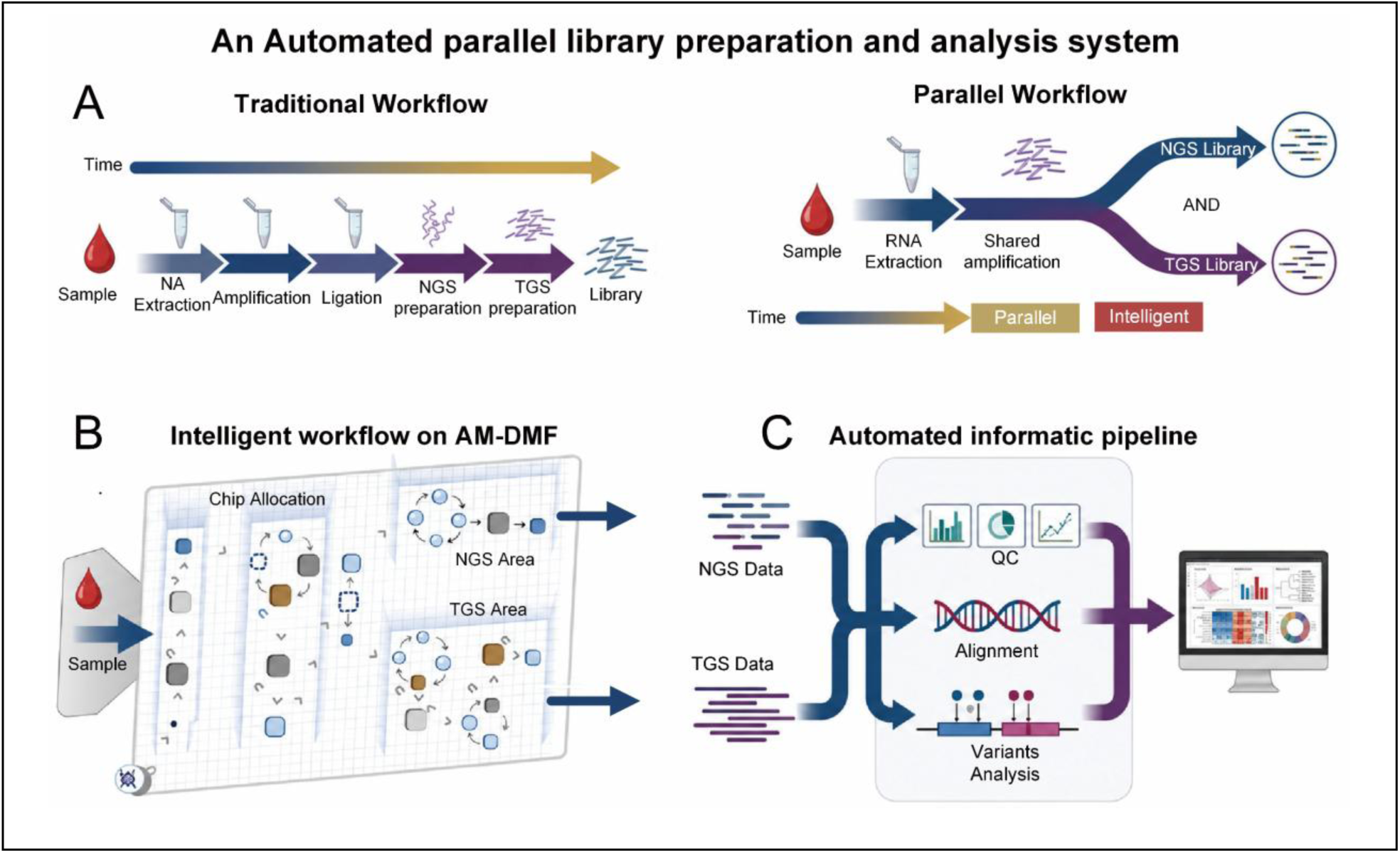
Workflow Overview. (A), Comparison between the traditional serial workflow (left) and the new parallel workflow (right). The conventional workflow performs RNA extraction, followed sequentially by NGS library preparation and TGS library preparation; the new workflow conducts shared amplification after RNA extraction, followed by parallel library preparation for both NGS and TGS platforms. (**B**), Intelligent sample allocation strategy on the integrated AM-DMF chip. (**C**), Automated bioinformatics pipeline capable of simultaneous analysis of NGS and TGS data.

Importantly, both the wet-lab microfluidics and the computational backend are intrinsically pathogen-agnostic: the system can be instantly reprogrammed for DENV, CHIKV, SARS-CoV-2, influenza, or any emerging threat simply by updating the multiplex primer panels and digital reference databases.

Using DENV and CHIKV as representative proof-of-concept targets—selected for their divergent genomic structures, expanding global prevalence, and urgent public health relevance—we demonstrate that the system reduces total reagent consumption by 72%, shortens library preparation time by 45%, and decreases the inter-batch coefficient of variation by 20% relative to conventional manual workflows. Clinical validation achieved 100% positive-percent agreement with RT-qPCR while simultaneously delivering high-resolution serotype, genotype, and quasi-species intelligence unobtainable by standard molecular diagnostics. By unifying real-time outbreak typing with deterministic genomic surveillance within an automated, field-deployable, and highly programmable ecosystem, this platform establishes a robust technological foundation for the next generation of viral genomic diagnostics.

## RESULTS

### Workflow of automated AM-DMF platform for short and long reads library preparation

To achieve fully automated processing of DENV from raw sample to sequencing library, we built the hardware and software system of an AM-DMF platform (Fig. 2A). The hardware integrated an electrode driving array, temperature control modules, illumination and imaging modules and a three-axis motion mechanism; the software enabled programmable control of droplet paths.

**Fig. 2.**
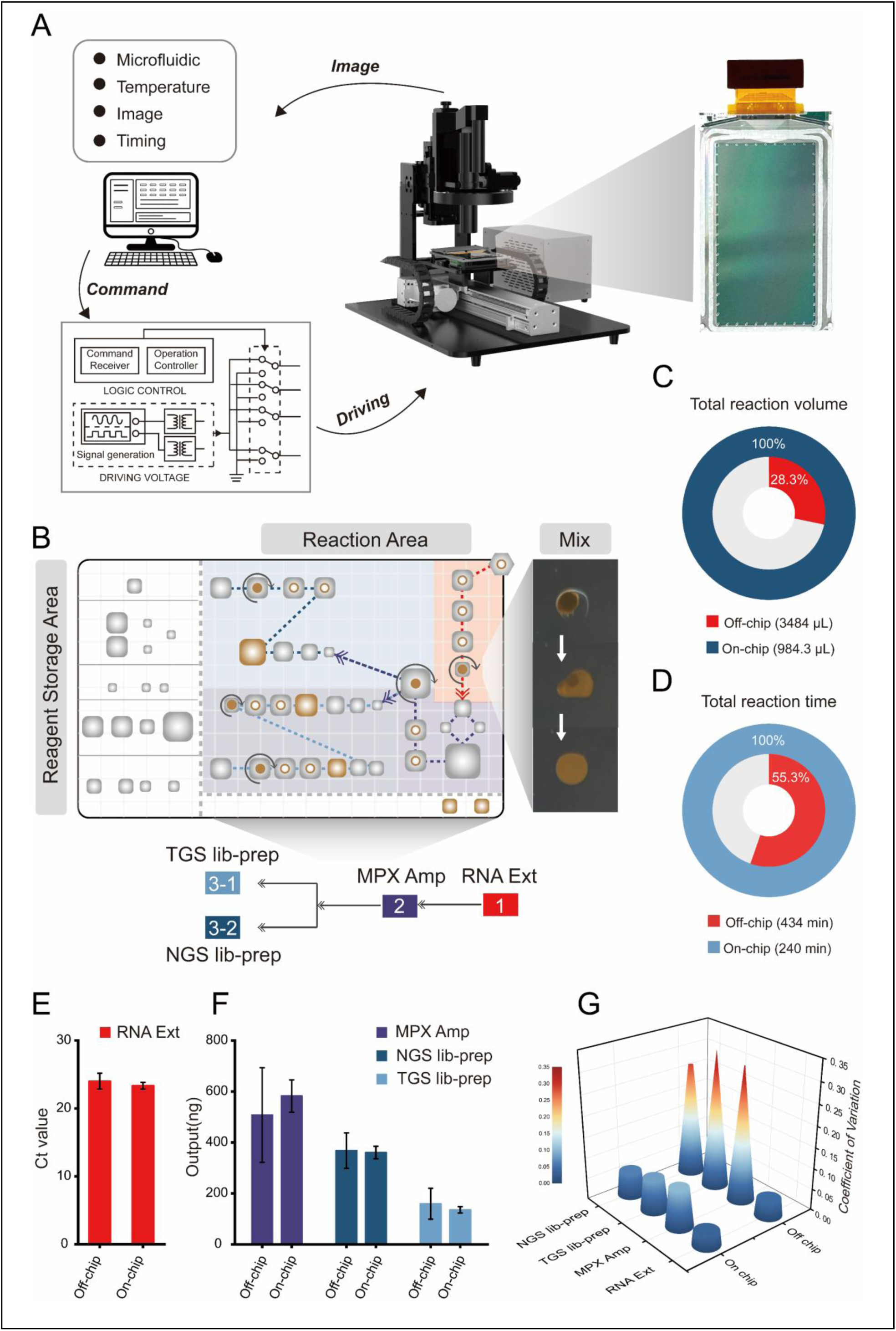
Optimization of AM-DMF platform for short and long reads library preparation. (A), AM-DMF integrated platform. (**B**), Schematic of the automated on-chip pre-processing workflow. (**C**), Comparison of total reagent consumption between off-chip and on-chip workflows. (**D**), Comparison of total processing time between off-chip and on-chip workflows. (**E**)-(**F**), Stepwise yield comparison between off-chip and on-chip workflows (extraction, multiplex amplification, TGS library preparation, NGS library preparation; n = 4). (**G**), stepwise inter-batch CV comparison between off-chip and on-chip workflows.

We designed an on-chip architecture suitable for the complex workflow (Fig. 2B and Movie S1). In Fig. 2B, the left side contained a 4 °C reagent storage area, and the right side was the reaction area, where steps were performed sequentially as extraction, multiplex amplification, NGS and TGS library preparation. To overcome the limitation of previous library preparation that could not handle raw samples because of restricted sample input and limited chip capacity, we integrated an external reservoir on chip (Supplementary Fig. S1A and B). This allowed raw samples to be directly loaded onto the chip, ensuring detection sensitivity while circumventing the volume limitation, thus achieving an automated “sample-in, library-out” process. Under software control, droplets moved, mixed and reacted according to set paths.

After optimizing the reaction systems and time parameters for each step (Supplementary Fig. S3, Supplementary Tables S2, S3), we compared the reagent consumption, total time and yields of the on-chip workflow with those of the manual procedure. The on-chip workflow reduced reagent consumption by 72% and shortened the total processing time by 45% (Fig. 2C–D). TGS provided a long read structural backbone, whereas NGS rigorously error corrected and quantified micro variant frequencies; both used the same sample aliquot, avoiding the sample degradation and time bias associated with separate library preparations. The yields of each step (extraction, multiplex amplification, TGS library preparation and NGS library preparation) were comparable to those of the manual procedure (Fig. 2E-F), indicating that automation did not compromise productivity.

We evaluated the inter-batch CV using four replicate experiments (Fig. 2G). Compared with manual operation, the on-chip platform reduced CV by 0.91% for extraction, 22.28% for multiplex amplification, 20.98% for TGS library preparation and 24.97% for NGS library preparation. These results demonstrate that the integrated AM-DMF platform not only saves reagents and shortens library preparation time, but also improves batch-to-batch consistency, providing a foundation for validation with clinical samples.

### Bioinformatics workflow for simultaneously analysis of sequencing data

Existing bioinformatics workflows are laborious for experimentalists, and the fragmented use of separate NGS and TGS pipelines further complicates true hybrid co-analysis.

Conventional bioinformatics relies on multiple independent tools and command line operations. Processing a single dataset typically requires 30–60 min and >15 commands, with manual batch management and high risk of errors, making it unfriendly to non-specialist personnel and unable to meet outbreak response demands (Fig. 3A).

**Fig. 3.**
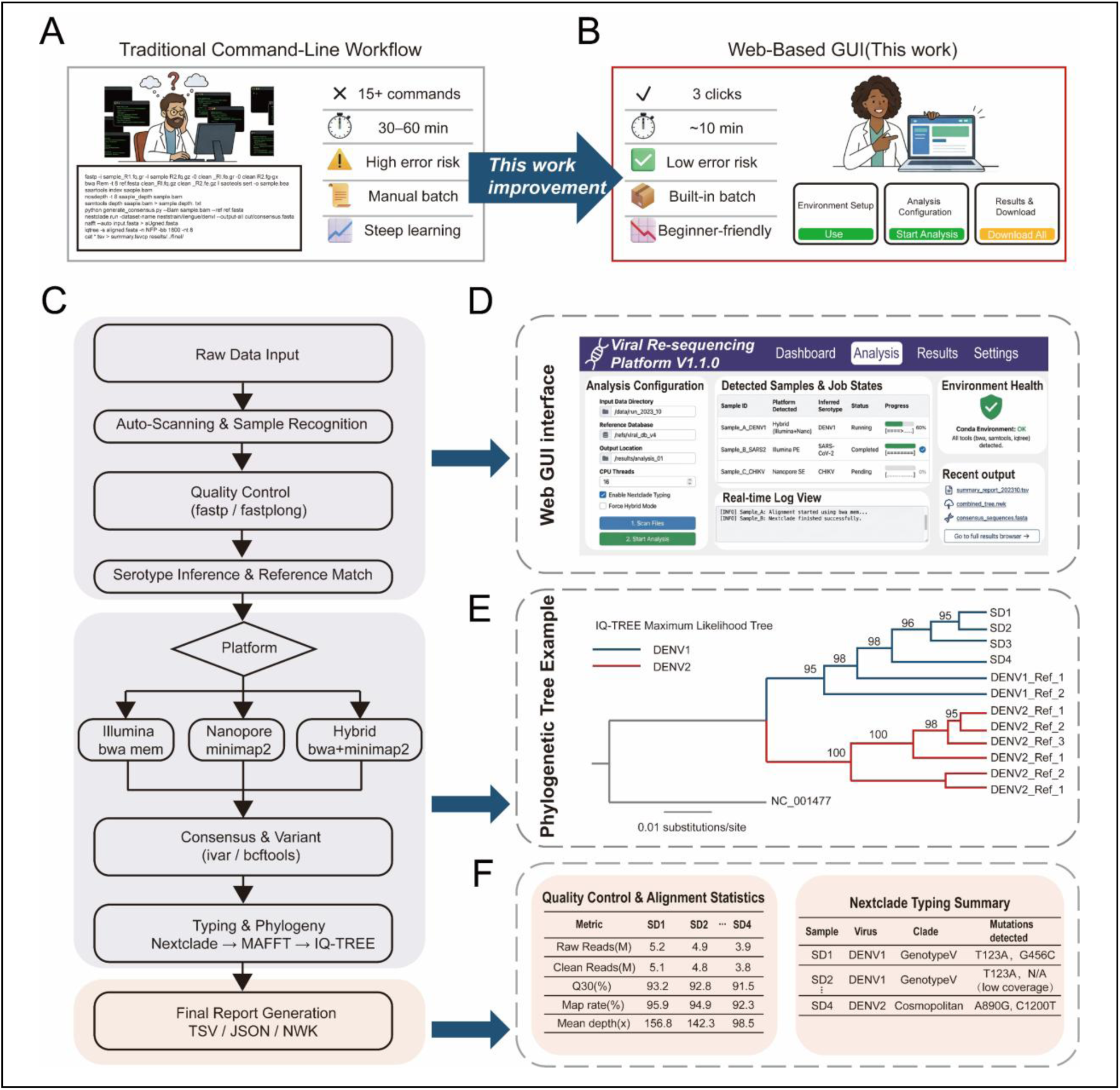
Workflow for bioinformatics analysis of sequencing data. (A), Conventional command-line analysis workflow. (B), Web-based GUI analysis workflow of this work. (C), Automated bioinformatics analysis pipeline. (D), Web-based GUI configuration and monitoring interface. (E), Example of phylogenetic tree output. f, Example of quality-control and typing result table.

To overcome this, we developed an automated workflow based on a web-based graphical user interface (GUI), designed to work with the AM-DMF pre-processing system (Fig. 3A, right). Users need only three clicks to complete analysis from raw data to a final visual report in ∼10 min. The workflow is compatible with both NGS and TGS data; the backend optimizes processing for each platform. Compared with command line, the workflow reduces analysis time by >70%, simplifies operations from >15 commands to three clicks, and lowers risk of human error (Fig. 3A, right).

The core innovation is its intelligent recursive scanning and modular logic (Fig. 3C). The system automatically executes: data integrity check, reference matching (recursive scanning), alignment, variant discovery, and report generation. Each module provides a visual interface: reference matching displays the most likely virus type (Fig. 3D); alignment shows read coverage depth; and the final report integrates virus identity, mutation spectrum, phylogeny (Fig. 3E) and QC metrics (Fig. 3F). The entire process requires no coding, achieving “drag and drop” automated analysis.

Using DENV clinical samples, the workflow reduced analysis time by >70% compared with traditional methods. In tests with mixed samples containing DENV1-4 serotypes, typing accuracy reached 100% (Fig. 3F). More importantly, the workflow simultaneously processes TGS (long read structural backbone) and NGS (error corrected micro variant frequencies) data, providing a complete information chain from viral identity to genomic architecture. This bioinformatics module integrates seamlessly with the AM-DMF wet experiment workflow, forming a field deployable sample to data solution.

### AM-DMF-sequencing system performance

We evaluated the performance of the AM-DMF-sequencing system for the four DENV serotypes (DENV1-4), including whole-genome coverage, library characteristics, limit of detection, batch-to-batch stability and cross-platform consistency.

Reference standards of DENV1-4 were first used for whole-genome sequencing. All serotypes achieved full genome coverage (Fig. 4A), indicating that the library preparation, primer design and analysis workflow were reliable. Library fragment lengths (Fig. 4B) showed that NGS libraries were centered at ∼150 bp and TGS libraries at ∼1200 bp, consistent with the technical characteristics of each platform. In low-concentration samples (≤10¹ copies/μL), some TGS fragments were shorter than expected, because of uneven amplification under low template amounts.

**Fig. 4.**
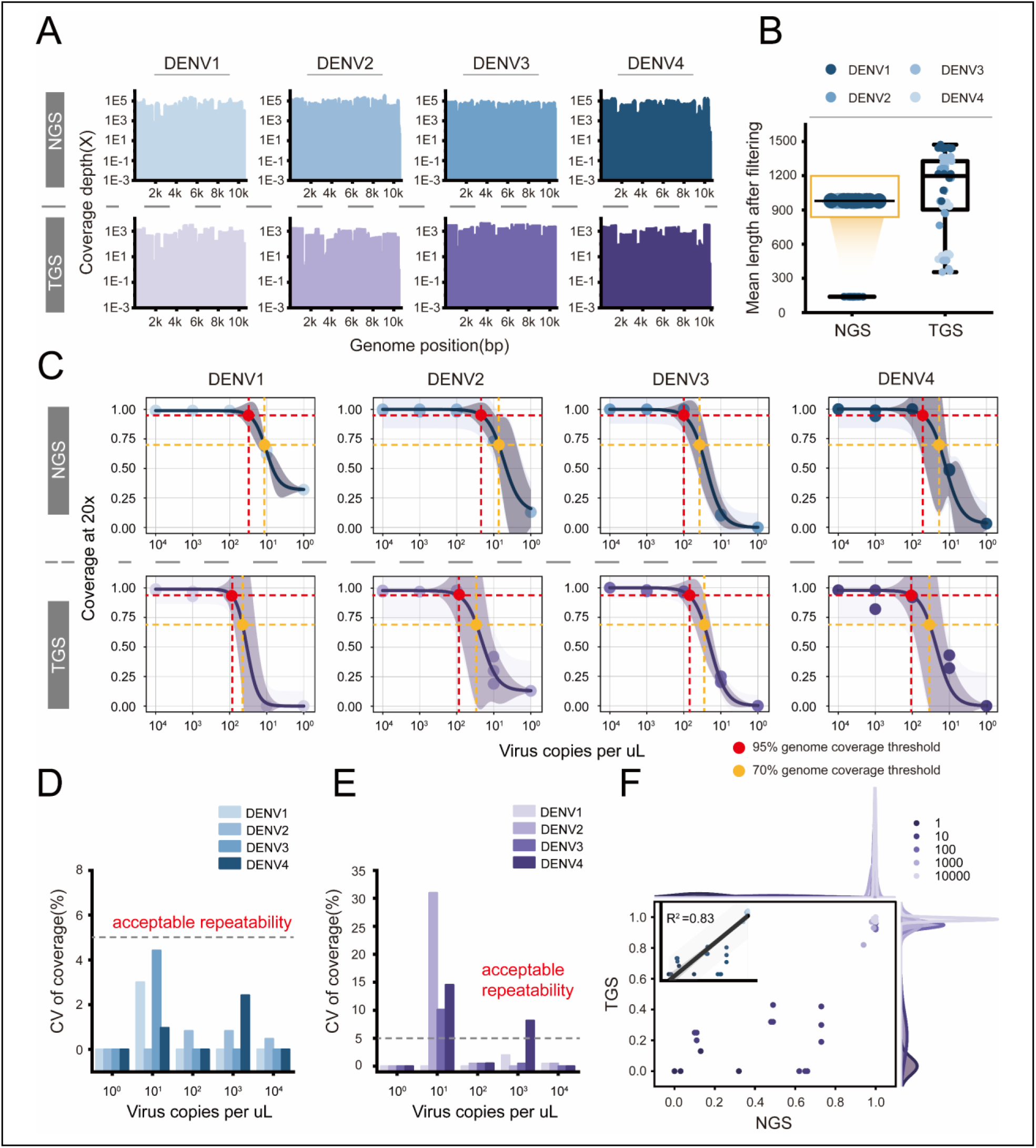
Performance evaluation of the arbovirus AM-DMF-sequencing system. (A), Whole-genome coverage depth of reference standards. (**B**), NGS and TGS library fragment length distribution. (**C**), Limit of detection for DENV1-4 by NGS and TGS (n = 3; 95% and 70% genome coverage threshold). (**D**), Coverage CV of NGS at different viral concentrations. (**E**), Coverage CV of TGS at different viral concentrations. (**F**), Correlation of coverage between NGS and TGS.

To quantify detection sensitivity, we adopted a scheme(*30*) and defined ≥70% coverage as the threshold for “successful sequencing”. We also recorded 95% coverage as a more stringent reference line (Fig. 4C). For DENV1-4, coverage was measured at different concentrations. The four serotypes reached 70% coverage at 10¹–10² copies/μL and 95% coverage at 10²–10³ copies/μL. TGS required slightly higher concentration for 95% coverage than NGS, so more samples fell into the 70–95% interval. The 70% threshold was sufficient for successful sequencing and phylogenetic construction, and we used it as the effective detection standard for clinical sample analysis.

The CV of coverage for three replicate experiments was calculated for both NGS and TGS across different concentration gradients (10⁰–10⁴ copies/μL) (Fig. 4D and Fig. 4E). NGS exhibited CV values <5% at all concentrations, indicating high intra-batch stability (according to CLSI guidelines, <5% is considered high precision)(*31, 32*). For TGS, at 10¹ copies/μL, the CV exceeded 5% for DENV2, DENV3 and DENV4, reflecting amplification and sampling fluctuations at low template amounts. At 10³ copies/μL, the CV for DENV4 also exceeded 5%, retrospective examination of the experimental records indicated that this outlier was caused by poor adapter ligation efficiency in a single library preparation and was not a systematic issue. At the lowest concentration (10⁰ copies/μL), the CV values of both platforms were very small because almost no data were detected, making the variation meaningless.

NGS-TGS correlation (Fig. 4F) showed that at ≥10² copies/μL, both platforms achieved high coverage with good consistency; at 10⁰–10¹ copies/μL, NGS coverage was higher than TGS, as NGS library preparation includes PCR amplification and generates more data than TGS. The coefficient of determination (R²) was 0.83, indicating strong correlation.

In summary, the AM-DMF-sequencing system achieved detection of DENV1-4, with a limit of detection (70% coverage) of 10¹–10² copies/μL and concordance between NGS and TGS results.

### Clinical validation with DENV samples

After reference standard evaluation, we validated the AM-DMF sequencing system using clinical samples. Ten RT-qPCR confirmed DENV positive samples were collected from a CDC, designated SD1–SD10, including four DENV1 and six DENV2 cases (Fig. 5B), with Ct values ranging from 16.64 to 30.31 (Fig. 5A).

**Fig. 5.**
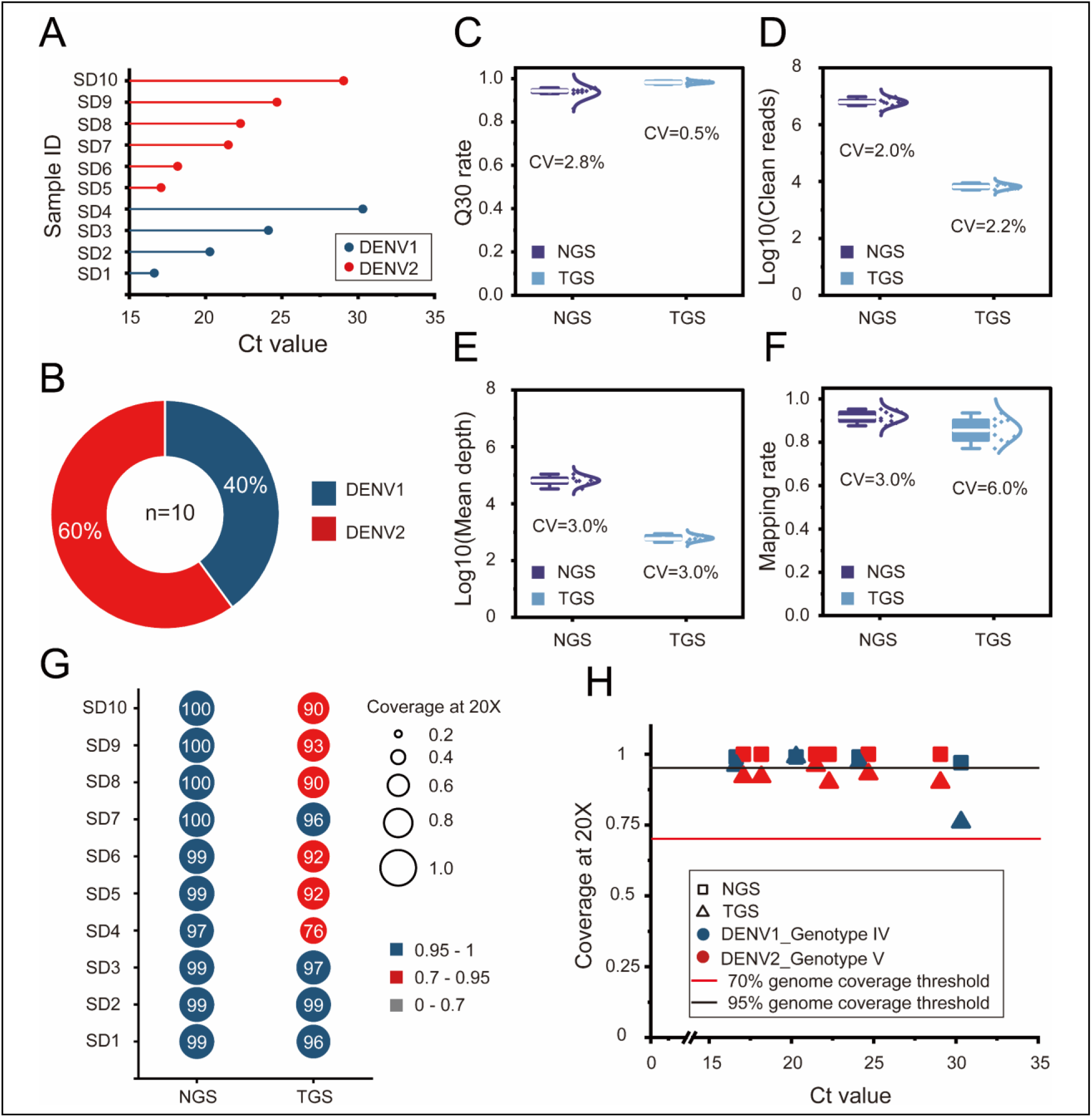
Clinical validation with DENV samples. (**A**), Distribution of clinical sample Ct values. (**B**), Clinical sample serotype proportion. (**C**)– (**F**), Variation of NGS and TGS sequencing quality metrics (Q30, clean reads, mean depth, mapping rate). (G), NGS and TGS genome coverage for each sample. (**H**), Relationship between Ct value and coverage (The 95% and 70% genome coverage threshold lines are indicated).

All samples were sequenced in parallel by NGS and TGS. Quality metrics (Q30, clean reads, mean depth and mapping rate) are shown as box plots (Fig. 5C–F). The inter batch CVs for NGS were 2.8%, 2.0%, 3.0% and 3.0%, respectively; for TGS: 0.5%, 2.2%, 3.0% and 6.0%. Both platforms showed good reproducibility, with the higher TGS CV for mapping rate (6.0%) consistent with earlier fluctuations at low concentrations.

Genome coverage of each sample is shown in Fig. 5G. NGS reached >95% coverage in all ten samples, demonstrating the robustness of the PCR amplification step in NGS library preparation for various sample types. TGS coverage was sample dependent: four samples (SD1, SD2, SD3, SD7) had >95% coverage; the remaining six (SD4, SD5, SD6, SD8, SD9 SD10) had 76%–93%(Supplementary Tables S4). The Ct values of these six spanned 17.08–30.31, and coverage differences were not explained by viral load alone (e.g., SD5, Ct=17.08, had 92% coverage; SD7, Ct=21.49, had >95%). Nevertheless, all TGS coverages exceeded the 70% threshold.

The relationship between Ct value and coverage is shown in Fig. 5H. All NGS points lay above the 95% line and showed no obvious correlation with Ct value. TGS points showed lower coverage with higher Ct value, and all points remained above the 70% line. One sample (SD7, Ct = 21.49, coverage >95%) exhibited slightly better tolerance than the trend would predict, further reflecting individual sample variation.

Using the built-in typing algorithm, all four DENV1 samples were identified as DENV1_Genotype IV, and all six DENV2 samples as DENV2_Genotype V, confirming that 70% coverage suffices for reliable phylogenetic analysis.

In the reference-standard DENV validation (Fig. 4), TGS achieved full genome coverage; in contrast, six clinical samples had 70–95% coverages. Because the same workflow was used, the difference likely originated from the samples themselves. TGS relies on long-fragment amplification, which is sensitive to RNA integrity11. Clinical samples may undergo RNA degradation during collection or storage, whereas the reference standard was cultured virus with highest integrity. Thus, batch-to-batch variation in RNA quality is the primary reason for the failure to achieve 100% TGS coverage.

In summary, the system achieved positive percent agreement in RT-qPCR-confirmed positive samples and provided richer genotypic information. NGS maintained high coverage across the full viral-load range, whereas TGS enabled effective serotyping (≥70% coverage) even with lower coverage. This difference indirectly validates the necessity of co-analysis.

### Clinical validation with CHIKV samples

The arbovirus family exhibits substantial genetic diversity. To assess the general applicability of the AM-DMF-sequencing system to different arboviruses, we performed a parallel validation using clinical CHIKV samples. Ten RT-qPCR-confirmed CHIKV-positive samples were collected from a CDC, with Ct values ranging from 11.78 to 35.29, covering a wide dynamic range from very high viral load to near the conventional detection limit (Fig. 6A).

**Fig. 6.**
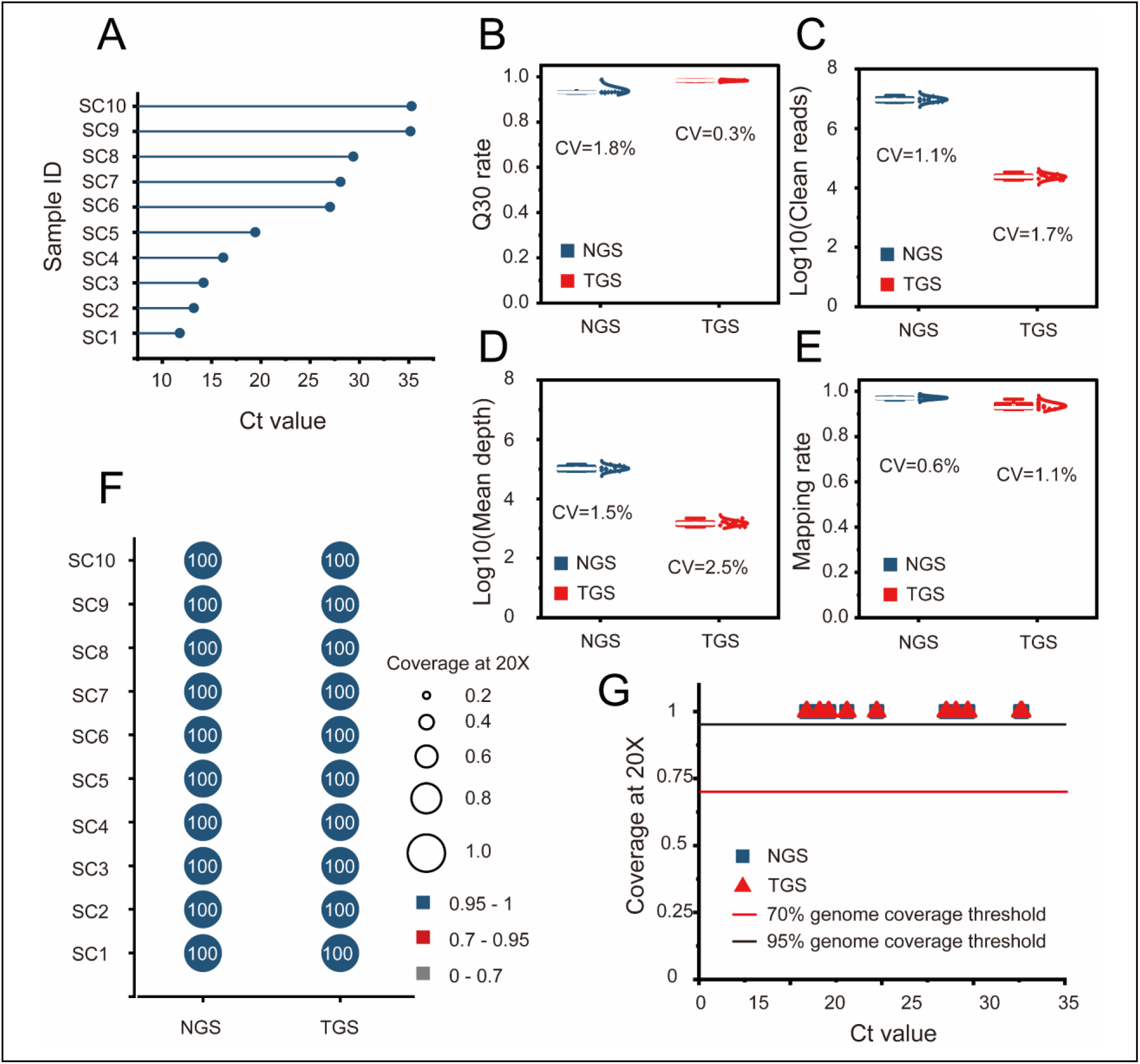
Clinical validation with CHIKV samples. (A), Distribution of clinical sample Ct values. (B)– (E), Variation of NGS and TGS sequencing quality metrics (Q30, clean reads, mean depth, mapping rate). (F), NGS and TGS genome coverage for each sample. (G), Relationship between Ct value and coverage (The 95% and 70% genome coverage threshold lines are indicated).

Box plots of sequencing quality metrics are shown in Fig. 6B–E. The inter-batch CVs for NGS were 1.8%, 1.1%, 1.5% and 0.6%, respectively; for TGS: 0.3%, 1.7%, 2.5% and 1.1%. Compared with DENV validation (NGS CV 2.0–3.0%, TGS CV 0.5–6.0%), the CHIKV data showed lower CV values, indicating better reproducibility.

All ten samples achieved 100% genome coverage for both NGS and TGS (Fig. 6F). Analysis of coverage vs. Ct value (Fig. 6G) showed that even for SC10, the sample with the highest Ct (35.29, very low viral load), NGS and TGS coverage remained at 100%, far above the 95% and 70% genome coverage threshold. This performance was markedly better than the DENV clinical validation, in which six samples had TGS coverages in the 70–95% range and none reached 100%.

Unlike the decreased TGS coverage observed for some DENV clinical samples, all CHIKV clinical samples achieved 100% genome coverage. Because the same AM-DMF platform and TGS workflow were used, and DENV reference standards had already shown that TGS can achieve full coverage when RNA is intact, the difference is most likely attributable to RNA quality of the clinical samples. TGS long-fragment amplification is highly dependent on template integrity. The CHIKV samples in this batch probably had better RNA integrity, allowing TGS to obtain complete coverage. This result confirms from the opposite perspective that RNA integrity is a key factor affecting TGS coverage and also demonstrates the high robustness of the platform when template quality is good.

In summary, the AM-DMF-sequencing system achieved 100% genome coverage for all CHIKV clinical samples(Supplementary Tables S5), with better reproducibility than for DENV. This validates the system’s versatility and highlights the influence of clinical sample RNA quality on TGS coverage. The platform holds good potential for field application during CHIKV and other arbovirus outbreaks.

## DISCUSSION

We have developed a complete sample-to-result system that enables, for the first time, strictly synchronous parallel library preparation for next generation sequencing (NGS) and third generation sequencing (TGS) directly from raw clinical samples, coupled with a bioinformatics pipeline. Compared with manual protocols, our AM-DMF platform reduces reagent consumption by 72%, shortens library preparation time by 45%, and lowers the inter batch CV by 20–25% for the most complex enzymatic steps. The accompanying pipeline analysis cuts time by >70% (from 30–60 min to ∼10 min) and reduces >15 command line operations to three clicks.

Using DENV1-4 reference standards, the system achieves a limit of detection (defined as ≥70% genome coverage) of 10¹-10² copies/μL. NGS and TGS complement each other: NGS provides robust coverage even at very low titers (≤10¹ copies/μL), whereas TGS offers long read structural information when viral load permits (≥10² copies/μL). In clinical validation with 20 RT-qPCR confirmed samples (10 DENV, Ct 16.6-30.3; 10 CHIKV, Ct 11.8-35.3), all samples were correctly serotyped and genotyped, confirming that 70% coverage is sufficient for reliable phylogenetic analysis.

The clinical data also reveal an important boundary condition: TGS coverage is highly sensitive to RNA integrity. All CHIKV samples achieved 100% TGS coverage, whereas some DENV clinical samples showed 76–93% coverage—still above the 70% threshold and sufficient for serotyping. Because DENV reference standards achieved full coverage using the same workflow, we attribute this difference primarily to batch-to-batch variation in RNA integrity.

Several limitations must be acknowledged. First, the sample size is modest (only 10 each for DENV and CHIKV), and validation is currently restricted to two arboviruses. Larger cohorts and broader viral panels (e.g., Zika, influenza, Ebola) are needed to establish generality. Second, the depth of co analysis between NGS and TGS remains limited; we have not yet fully exploited their synergistic potential for haplotype phasing and recombination detection. Future work will include more sophisticated multi modal fusion algorithms. Third, the current platform does not integrate the sequencing step itself; sequencing still requires a separate instrument (e.g., Nanopore). Fourth, no negative controls were included, so specificity remains to be formally evaluated.

Looking forward, we aim to expand viral coverage, conduct large scale prospective clinical studies, and deepen the co analysis framework. Most importantly, we will attempt to integrate a miniature sequencer onto the AM-DMF chip, achieving a fully enclosed, end to end automated workflow from sample input to genomic report. With further cost reduction and standardization, this platform holds promise as a routine tool for arbovirus outbreak surveillance and molecular diagnostics in resource limited settings.

In summary, the integrated AM-DMF sequencing and bioinformatics system reported here effectively addresses multiple bottlenecks from sample-to-result. Its automation, miniaturization and dual platform compatibility lay a solid technical foundation for the next generation of viral genomic diagnostics.

## MATERIALS AND METHODS

### Active-matrix DMF chip

The AM-DMF chip used in this work had a sandwich-like configuration, with the top and bottom glass plates separated by a 400 μm-thick double-sided adhesive gasket. The top plate contained a reagent injection hole, and the bottom plate was a 0.5 mm-thick glass substrate. An active electrode matrix was fabricated on the bottom plate using a standard a-Si:H thin-film transistor (TFT) process (manufactured by TIANMA Microelectronics, Shenzhen, China). The matrix consisted of 720 rows and 1,280 columns, providing 921,600 independent TFT electrodes with a single-pixel size of 70 μm (Supplementary Fig. S1C,D). Chip assembly was carried out by ACX instruments (Cambridge, UK) and ACXEL Micro&Nano Tech (Foshan, China).

A custom-designed integrated circuit (IC) was used to drive the chip. Timing and data signals were generated by a high-speed microcontroller and transmitted to the IC (Supplementary Fig. S1E,F). After receiving a small number of input signals, the IC output row and column addressing signals through internal logic operations, thereby achieving independent control of each electrode. Compared with our previous work23, the present design integrated more electrodes on the same chip area, reduced the single-pixel size to 70 μm and required only 40 connecting wires to drive the chip, improving performance while reducing driving complexity.

To enable direct loading of raw samples, an external reservoir was integrated on the chip (Supplementary Fig. S1B, attached to a 1 mm hole on the chip via adhesive bonding). This reservoir accommodated large-volume samples and, through pre-set droplet paths, introduced the sample into the reaction region, facilitating a fully automated “sample-in, library-out” process.

### AM-DMF cartridge and system

Precise temperature control in the range of 4 °C–98 °C was required for reagent storage and pre-processing reactions. We therefore adopted a temperature control component from our previous work(*33*), which combined TEC heating with fuzzy-PID principles to achieve rapid on-chip temperature ramping and constant-temperature maintenance. To enable independent temperature management of different functional zones, three independent temperature-controlled regions were designed on the chip, corresponding to the reagent storage area, the NGS reaction area and the TGS reaction area.

A system integrating chip driving, chip imaging, interactive software and temperature control was developed for automated library preparation. The temperature control module and the chip driving module were mounted together on a set of guide rails; by moving the rails, the system could switch to bright-field imaging mode, and a high-definition camera (Hikvision, China) was used to monitor the chip status in real time. Detailed system configuration is shown in Supplementary Fig. S2. The custom system supported automatic reagent addition through the chip driving module, automatic execution of temperature control programs (with independent regulation of each zone) and complete recording of the entire reaction process.

### Samples and reference standards

DENV reference standards (laboratory-cultured virus strains) were cultured in BSL-2 labs of Guangdong Second Provincial General Hospital of Jinan University. Clinical specimens were confirmed by RT-qPCR and collected from the Affiliated Guangdong Second Provincial General Hospital of Jinan University. They included 10 DENV-positive samples (4 DENV1 and 6 DENV2) and 10 CHIKV-positive samples (all CHIKV). Sample collection was performed under institutional ethics committee-approved protocol (W96-027M-202502), and all samples were identified. Specimens were stored at –80 °C and transported on dry ice.

### Chemicals and sample pre-processing

Detailed information on the reagents used in this work is listed in Supplementary Table S1. A 100 μL liquid sample (reference standard or clinical plasma) was mixed with 300 μL lysis buffer and gently inverted 3–5 times. An equal volume (400 μL) of binding buffer was then added and again gently inverted 3–5 times. Finally, 20 μL of BeyoMag magnetic beads were added, giving a total volume of 820 μL. This mixture was directly used for on-chip extraction.

### On-chip wet-experiment pre-processing workflow

First, the required reaction reagents were pipetted into the chip through the injection port and delivered to the corresponding 4 °C reagent storage area according to a pre-set program. The prepared sample mixture (820 μL) was then added to the external reservoir of the chip. The chip automatically executed the pre-set program, sequentially performing magnetic bead mixing, washing and RNA elution. Reverse transcription, multiplex amplification, NGS library preparation and TGS library preparation were all carried out automatically in the pre-programmed order (see Movie S1 and Supplementary Table S3 for details). The final products were pipetted out of the chip, quantified using a Qubit 4 fluorometer (Thermo Fisher) and stored at –80 °C for subsequent sequencing.

### Sequencing

NGS libraries were sequenced on an Illumina NovaSeq 6000 platform in PE150 paired-end mode with a target data volume of 1 Gb per sample. TGS libraries were sequenced on a GENEUS G-seq500 platform (equipped with a G-MK02 microfluidic chip) for 120 min.

### System and time optimization

To adapt the reactions to on-chip automated operation, we optimized the volumes of the multiplex amplification, NGS library preparation and TGS library preparation reactions. The multiplex amplification system was reduced to 1/2 of its original volume, and the NGS library preparation system was reduced to 1/10 of its original volume (the TGS system used the same 1/10 volume). The reduced volumes still produced yields sufficient for downstream sequencing (Supplementary Fig. S3B, C). On-chip washing and elution volumes were optimized to 7 μL and 9 μL, respectively, both reaching plateau yields (Supplementary Fig. S3B, E). For time reduction, we shortened the duration of each reaction step and replaced the conventional stepwise operation with a concurrent mixing-while-reacting strategy. These changes reduced the total on-chip process time by 45% compared with the manual procedure (Supplementary Table S2–S3, Supplementary Fig. S3).

### Bioinformatics analysis

Sequencing data (FASTQ format) were fed into a self-developed GUI pipeline (based on the Python/Flask framework). The pipeline was compatible with both NGS and TGS data and automatically performed the following steps: (1) quality control (fastp for NGS, fastplong for TGS); (2) intelligent reference database matching (recursive scanning of DENV1-4 and CHIKV reference genomes); (3) sequence alignment (BWA-MEM for NGS, minimap2 for TGS); (4) variant calling and consensus sequence generation (using iVar and samtools); and (5) report generation that includes coverage curves, mutation spectra, phylogenetic trees and serotype/genotype identification (Supplementary report).

### Performance evaluation

Total time and CV. The total reagent consumption and total processing time (from sample input to library output) of the on-chip workflow was compared with those of the manual procedure; details are shown in Fig. 2C–D and Supplementary Table S3. The inter-batch CV for each step was calculated from four replicate experiments (standard deviation/mean × 100%).

Limit of detection DENV1-4 reference standards were serially diluted ten-fold (10⁰–10⁴ copies/μL) and each concentration was tested in triplicate. A 95% genome coverage probability was defined as high confidence detection, and a 70% genome coverage probability was defined as successful sequencing.

### Statistical analysis

All experimental data are presented as mean ± standard deviation. The CV was calculated as (standard deviation/mean) × 100%. The limit of detection was determined by logistic non-linear fitting. The correlation between NGS and TGS coverage was assessed using the Pearson correlation coefficient (R²). Positive percent agreement was calculated as described above.

## Acknowledgments

**Funding**

This work was supported by the National Key R&D Program of China (No. 2023YFF0721500), the National Natural Science Foundation of China (No. T2541056), the Suzhou Basic Research Project (No. SSD2023013), and the Guangdong Scientific and Technological Project (No. 2025A0505010003).

**Author contributions**

B. Zhang and Z. Fang conceived the concept and designed the experiments. B. Zhang, Q. Liu and J. Ji developed the platforms and software. B. Zhang, S. Hu and Y. Wang performed the research. B. Zhang, C. Yong, X. Lai, Y. Feng and J. Li(Jiahao Li) analyzed the data. B. Zhang, J. Yu, C. Jiang and H. Ma wrote the paper. A. Nathan, J. Li(Jinhua Li), C. Yu and H. Ma provided professional advice and supervised the project. All authors reviewed and approved the final manuscript.

**Competing interests**

The authors declare no conflict of interest..

**Data, code, and materials availability:** The data that support the findings of this study are available from the corresponding author upon reasonable request.

