## Supplementary Materials for "An active-matrix digital microfluidic platform for simultaneous short- and long-read viral genomic surveillance"

Science Advances

AAAS

**This PDF file includes:**

Figs. S1 to S3
Tables S1 to S5

**Other Supplementary Materials for this manuscript include the following:**

Movies S1
Source code S1
Report S1

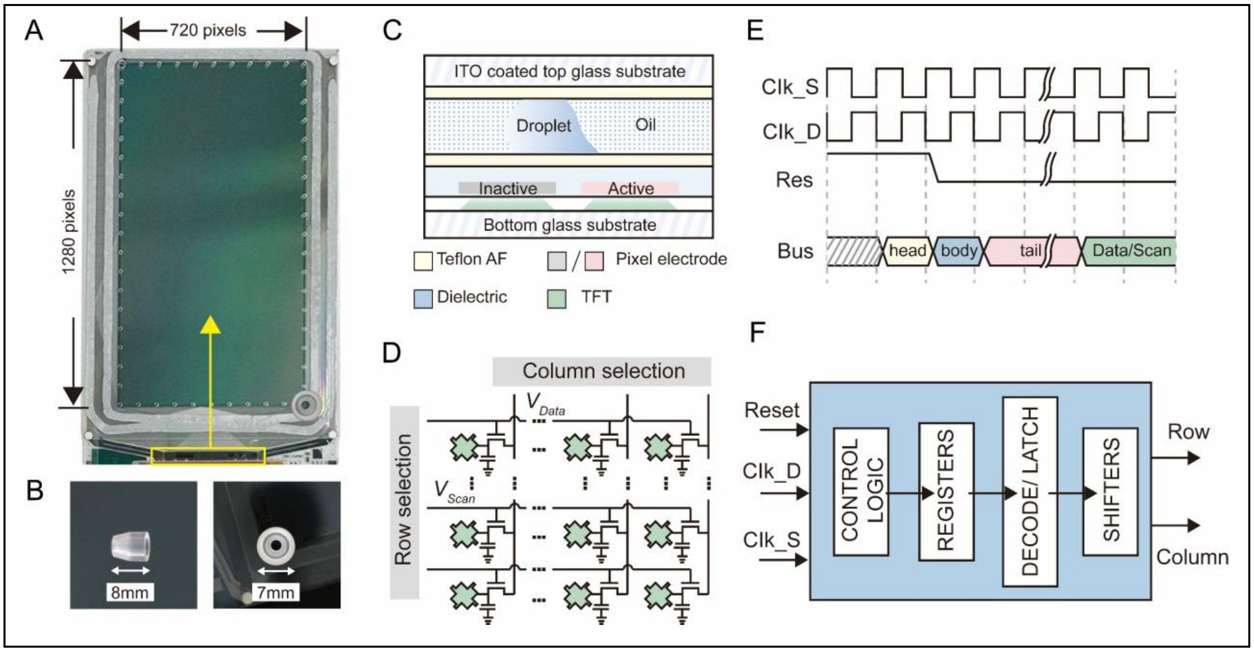


**Fig. S1.**
**AM-DMF chip and electronic integrated circuit.** a, Photograph of the AM-DMF chip with IC and external reservoir. b, External reservoir for large-volume sample extraction. c, Layer structure diagram of the chip. d, Peripheral driving circuit of the integrated circuit. e, Typical timing diagram of the IC driver. f, IC design architecture (control logic, registers, decode/latch, level shifter).

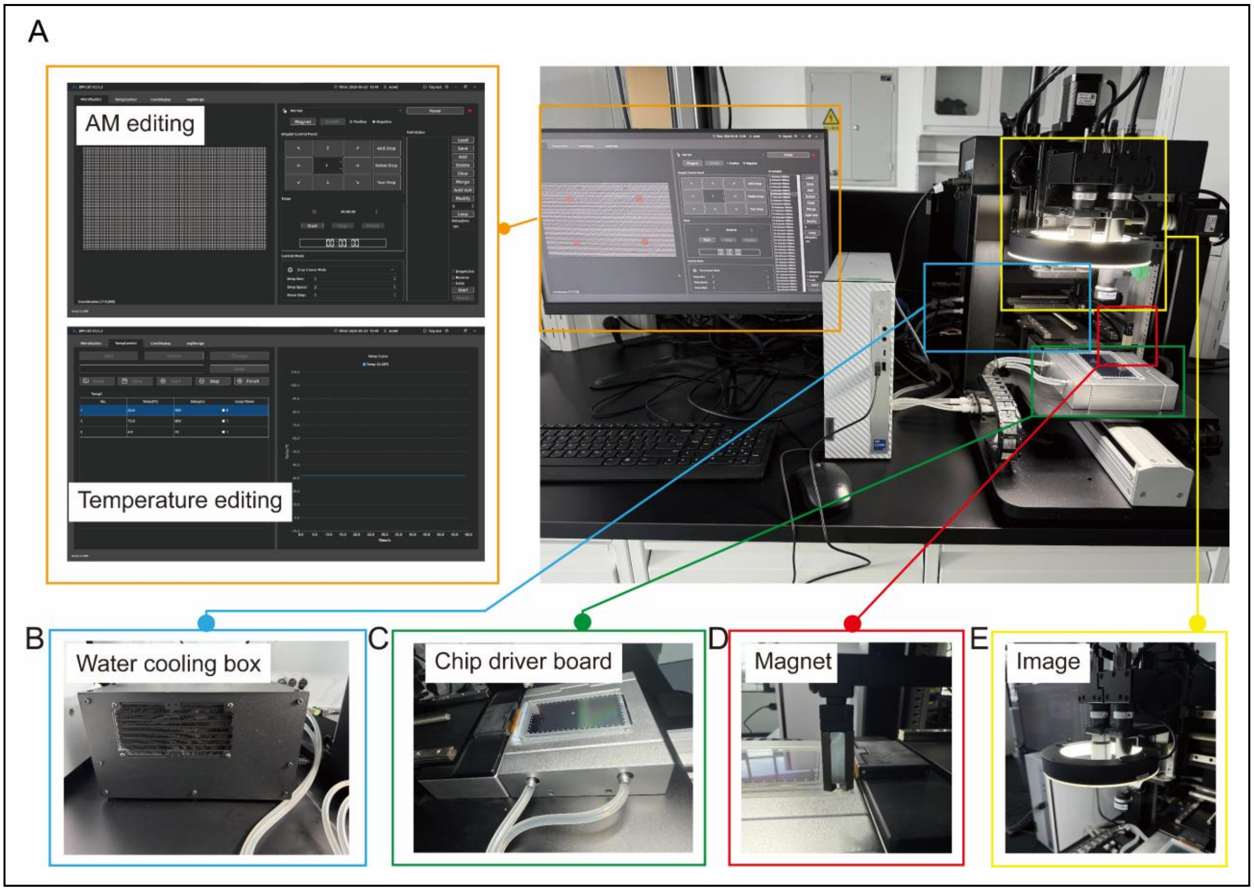


**Fig. S2.**
**System developed for AM-DMF sequencing pre-processing, consisting of the following components:** a, Interactive software for editing and operating droplet paths, real-time display of droplet movement, and temperature setting with temperature-curve display. b, Water-cooling box for TEC heat dissipation. c, Temperature control module and chip driving module mounted together. d, Magnetic function using a magnetic lens design to enhance magnetic field strength and flux focusing. e, High-speed camera and lens forming the chip imaging module for displaying chip view and real-time observation of droplet paths.

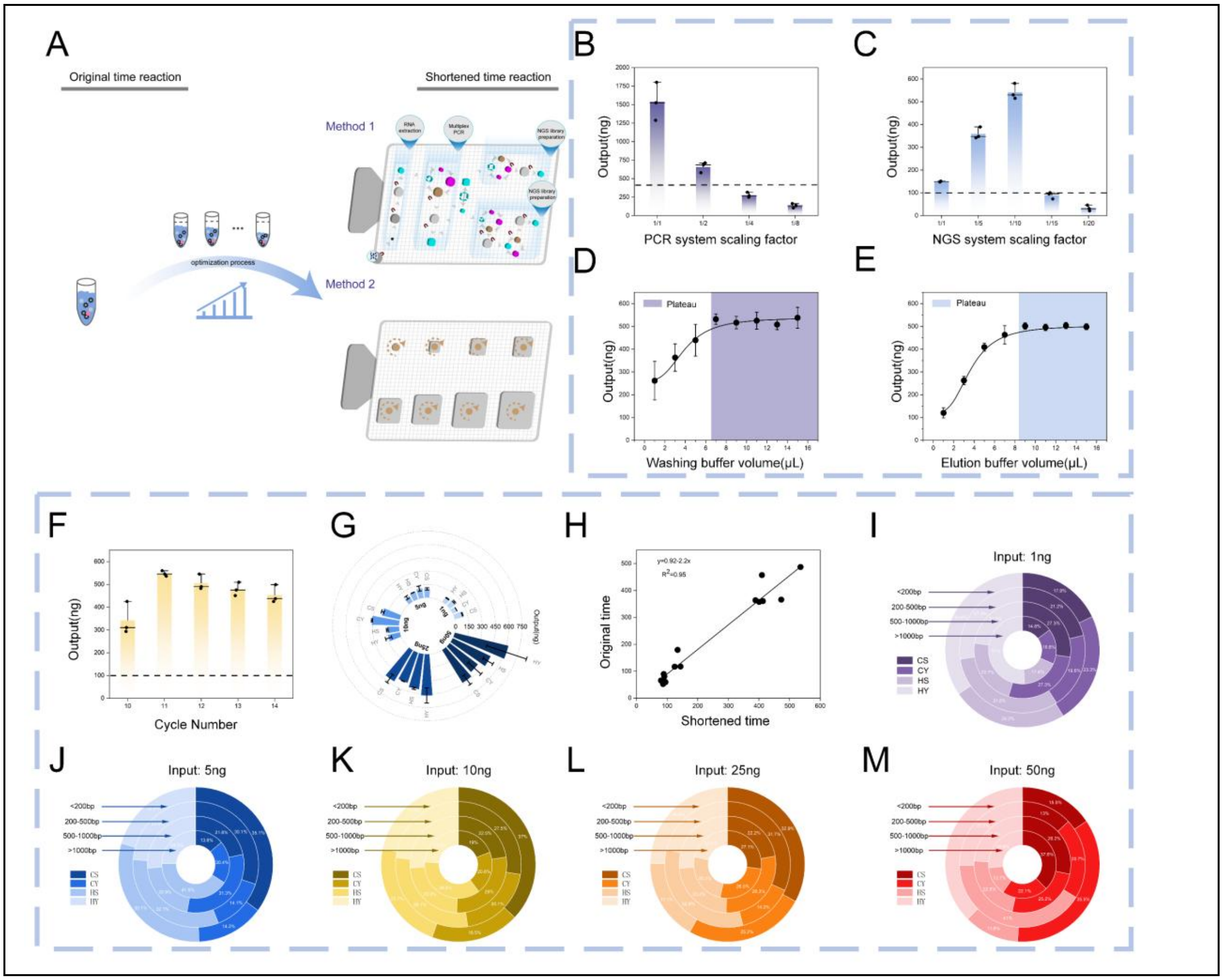


**Fig. S3.**

**Optimization of the on-chip pre-processing workflow on the AM-DMF platform.** a, Schematic of the optimization process converting the original reaction system (large volume, long time) to an on-chip automated, miniaturised, shortened workflow. Volumes were first optimized (b–c): the multiplex amplification system was reduced to 1/2 of its original volume, and the NGS library preparation system to 1/10, both still meeting experimental requirements; TGS used the same 1/10 system to maintain volume consistency. d, Optimization of on-chip washing and elution volumes: 7 μL for washing and 9 μL for elution reached plateau yields. Reaction time was then optimized (f): at 1 ng DNA input, the optimal number of NGS library preparation cycles was 11, giving the highest yield.

g, Comparison of NGS library yields under on-chip shortened time, on-chip original time, manual shortened time and manual original time (n = 3). h, Consistency of NGS library yields between shortened and original time conditions ($R^2 = 0.95$). i–m, Fragment size distribution of libraries prepared under on-chip shortened time, on-chip original time, manual shortened time and manual original time with different DNA inputs (1, 5, 10, 25, 50 ng).

**Table S1. Chemicals used in this work**

| Reagent | Company | Experiment |
|---|---|---|
| Magnetic bead-based viral DNA/RNA extraction kit | Beyotime (Shanghai, China) | RNA extraction |
| Carrier RNA | Beyotime (Shanghai, China) | RNA extraction |
| Dengue virus serotyping real-time PCR kit | IANLONG (China) | qPCR quantification |
| Dengue virus whole-genome capture kit | BAIYITECH (China) | Reverse transcription and amplification |
| LunaScript RT SuperMix Kit | New England Biolabs(USA) | Reverse transcription |
| Q5 Hot Start High-Fidelity 2X Master Mix | New England Biolabs(USA) | Amplification |
| FS Pro DNA Library Prep Kit V2 | ABclonal (China) | NGS library preparation |
| Universal Sample Preparation Kit for Nanopore Sequencing | GENEUS (China) | TGS library preparation |
| Qubit 1X dsDNA HS/BR Assay Kits | Thermo Fisher (USA) | DNA quantification |
| Nuclease-free water | Thermo Fisher (USA) | Elution |
| 10% Pluronic F68 solution | Sigma-Aldrich (USA) | Whole process |
| Silicone oil (2 cSt) | Sigma-Aldrich (USA) | Whole process |
| Teflon-AF | Chemours (USA) | Chip assemble |

**Table S2. Comparison of original and optimized reaction conditions for NGS library preparation.**

| Step | Reaction condition (Original) | Reaction condition (Optimized) |
|---|---|---|
| DNA | 1/5/10/25/50 ng | 1/5/10/25/50 ng |
| Fragmentation & end repair | 32°C-5 min; 72°C-30 min | 32°C-5 min; 72°C-15 min |
| Adapter ligation | 22°C-15 min | 22°C-10 min |
| Purification | 30 min | 13 min |
| PCR | 98°C-1 min; (98°C-10 s, 60°C-30 s, 72°C-30 s) 6-10 cycles; 72°C-1 min | 98°C-1 min; (98°C-5 s, 60°C-15 s, 72°C-15 s) 6-10 cycles; 72°C-1 min |
| Purification | 30 min | 13 in |

- The pre treatment workflow adopts a strategy of simultaneous mixing and reaction, which shortens the overall process time.

- The purification times indicated apply to each step of the pre treatment procedure.

Reaction time optimization for multiplex amplification and TGS library preparation was not achieved; only the mixing while reacting strategy and purification steps were optimized.

**Table S3. Comparison of original and optimized reaction conditions for NGS library preparation.**

| Step | Procedure | Item | Reagents | Volume Off-chip(uL) | Volume On-chip(uL) | Time Off-chip (min) | Time On-chip (min) |
|---|---|---|---|---|---|---|---|
| 1 | RNA extraction | Sample preparation | Sample | 820 | 820 | 5 | 5 |
| 2 | | | Lysis buffer | | | | |
| 3 | | | Binding buffer | | | | |
| 4 | | | Magnetic beads | | | | |
| 5 | | | Carrier RNA | | | | |
| 6 | | Wash | Wash buffer 1 | 600 | 7 | 30 | 13 |
| 7 | | | Wash buffer 2 (×2) | 600 | 7 | | |
| 8 | | Elution | Elution buffer | 50 | 6 | | |
| 9 | Reverse transcription | Reverse transcription | RT super mix | 3 | 2 | 27 | 26 |
| 10 | | | RNA | 12 | 6 | | |
| 11 | Whole genome amplification | Whole genome amplification | cDNA | 2.5 | 1.25 | 125 | 114 |
| 12 | | | Hotstart hifi mix | 12.5 | 6.25 | | |
| 13 | | | Nuclease-free water | 6 | 3 | | |
| 14 | | | Primer pool A/B | 4/4 | 2/2 | | |
| 15 | | Purification | Purification beads | 30 | 15 | 30 | 13 |
| 16 | | Wash | 80% ethanol (×2) | 200 | 7 | | |
| 17 | | Elution | Nuclease-free water | 50 | 15 | | |
| 18 | NGS library preparation | Fragmentation & end repair | DNA | 32 | 3.2 | 40 | 20 |
| 19 | | | FS pro buffer I | 5 | 0.5 | | |

| | | | | | | | |
|---|---|---|---|---|---|---|---|
| 20 | | | FS pro enzymes II | 13 | 1.3 | | |
| 21 | | Adapter ligation | End-repaired product | 50 | 5 | 20 | 10 |
| 22 | | | FS pro ligation buffer II | 20 | 2 | | |
| 23 | | | Ligase enzymes | 5 | 0.5 | | |
| 24 | | | Working adapter | 5 | 1 | | |
| 25 | | Purification | Purification beads | 64 | 6.8 | 30 | 13 |
| 26 | | Wash | 80% ethanol (×2) | 200 | 7 | | |
| 27 | | Elution | Nuclease-free water | 21 | 2.5 | | |
| 28 | | PCR | Adapter-ligated product | 20 | 2.5 | 18 | 8 |
| 29 | | | 2X PCR mix | 25 | 2.5 | | |
| 30 | | | UDI primer | 5 | 0.5 | | |
| 31 | | Purification | Purification beads | 50 | 5 | 30 | 13 |
| 32 | | Wash | 80% ethanol (×2) | 200 | 7 | | |
| 33 | | Elution | Nuclease-free water | 31 | 9 | | |
| 34 | TGS library preparation | DNA repair & ligation | DNA | 35 | 3.5 | 14 | 9 |
| 35 | | | Reaction solution R-1 | 10 | 1 | | |
| 36 | | | Adapter | 5 | 0.5 | | |
| 37 | | Post-ligation | Adapter-ligated product | 25 | 5 | 35 | 30 |
| 38 | | | Reaction solution R-2 | 5 | 0.5 | | |

| | | | | | | | |
|---|---|---|---|---|---|---|---|
| 39 | | Purification | Purification beads | 60 | 6 | | |
| 40 | | Wash | LW | 200 | 7 | 30 | 13 |
| 41 | | Elution | Nuclease-free water | 12 | 9 | | |
| | total | | | 3484 | 984.3 | 434 | 240 |

- Off chip total time: 7.5 h (450 min); on chip total time: 3.9 h (235 min, based on the longer NGS protocol, since NGS and TGS are performed simultaneously on chip). Time saving: 45%.
- Off chip total reagent volume: 3443.5 μL; on chip total reagent volume: 982.7 μL. Reagent saving: 72%.

**Table S4. NGS characteristics of the 20 plasma samples used in this work**

| Sample no. | Serotype | Genotype | Ct Value | Q30(%) | Total reads | Mean depth | mapping rate | Genome coverage(%) |
|---|---|---|---|---|---|---|---|---|
| SD1 | DENV1 | Genotype IV | 16.64 | 94.74 | 7133882 | 78272.11 | 89.99 | 99 |
| SD2 | DENV1 | Genotype IV | 20.28 | 94.55 | 5758744 | 62599.96 | 89.77 | 99 |
| SD3 | DENV1 | Genotype IV | 24.12 | 93.14 | 6199592 | 61007.3 | 87.62 | 99 |
| SD4 | DENV1 | Genotype IV | 30.31 | 86.23 | 2982400 | 32904.75 | 91.02 | 97 |
| SD5 | DENV2 | Genotype V | 17.08 | 94.82 | 8895186 | 97292.53 | 93.85 | 100 |
| SD6 | DENV2 | Genotype V | 18.17 | 94.30 | 4736840 | 49133.45 | 92.21 | 100 |
| SD7 | DENV2 | Genotype V | 21.49 | 93.71 | 9574678 | 108921.55 | 94.99 | 100 |
| SD8 | DENV2 | Genotype V | 22.29 | 95.60 | 6240778 | 65699.97 | 93.69 | 100 |
| SD9 | DENV2 | Genotype V | 24.68 | 93.74 | 5594202 | 53049.34 | 88.98 | 100 |
| SD10 | DENV2 | Genotype V | 29.05 | 95.78 | 6698220 | 74233.45 | 95.3 | 100 |
| SC1 | CHIKV | II-ECSA | 11.78 | 93.26 | 10079794 | 112371 | 96.14 | 100 |
| SC2 | CHIKV | II-ECSA | 13.2 | 92.97 | 8209638 | 91924.66 | 96.93 | 100 |
| SC3 | CHIKV | II-ECSA | 14.19 | 93.23 | 9231596 | 104166.83 | 97.37 | 100 |
| SC4 | CHIKV | II-ECSA | 16.19 | 93.18 | 12953354 | 144755.19 | 96.72 | 100 |
| SC5 | CHIKV | II-ECSA | 19.44 | 92.74 | 8645412 | 96962.63 | 97.13 | 100 |
| SC6 | CHIKV | II-ECSA | 27.03 | 93.20 | 10048474 | 112338.48 | 96.55 | 100 |
| SC7 | CHIKV | II-ECSA | 28.09 | 93.37 | 12174894 | 135495.67 | 96.99 | 100 |
| SC8 | CHIKV | II-ECSA | 29.38 | 93.04 | 7523312 | 85770.23 | 98.59 | 100 |
| SC9 | CHIKV | II-ECSA | 35.18 | 93.10 | 7726902 | 87150.75 | 96.99 | 100 |
| SC10 | CHIKV | II-ECSA | 35.29 | 98.63 | 7943238 | 85626.15 | 97.06 | 100 |

**Table S5. TGS characteristics of the 20 plasma samples used in this work**

| Sample no. | Serotype | Genotype | Ct Value | Q30(%) | Total reads | Mean depth | mapping rate | Genome coverage(%) |
|---|---|---|---|---|---|---|---|---|
| SD1 | DENV1 | Genotype IV | 16.64 | 98.11 | 5071 | 439.98 | 90.74 | 96 |
| SD2 | DENV1 | Genotype IV | 20.28 | 97.45 | 8912 | 877.94 | 77.14 | 99 |
| SD3 | DENV1 | Genotype IV | 24.12 | 98.70 | 7178 | 551.46 | 93.53 | 97 |
| SD4 | DENV1 | Genotype IV | 30.31 | 97.54 | 8571 | 583.18 | 83.42 | 76 |
| SD5 | DENV2 | Genotype V | 17.08 | 98.92 | 7807 | 751.52 | 87.62 | 92 |
| SD6 | DENV2 | Genotype V | 18.17 | 98.08 | 5059 | 538.88 | 81.03 | 92 |
| SD7 | DENV2 | Genotype V | 21.49 | 97.77 | 6928 | 765.69 | 89.44 | 96 |
| SD8 | DENV2 | Genotype V | 22.29 | 98.84 | 6214 | 560.3 | 90.23 | 90 |
| SD9 | DENV2 | Genotype V | 24.68 | 98.48 | 5731 | 592.94 | 80.09 | 93 |
| SD10 | DENV2 | Genotype V | 29.05 | 98.76 | 5707 | 570.93 | 82.65 | 90 |
| SC1 | CHIKV | II-ECSA | 11.78 | 98.52 | 20307 | 1277.27 | 92.79 | 100 |
| SC2 | CHIKV | II-ECSA | 13.2 | 98.53 | 26911 | 1692.96 | 93.89 | 100 |
| SC3 | CHIKV | II-ECSA | 14.19 | 98.45 | 26229 | 1679.62 | 92.79 | 100 |
| SC4 | CHIKV | II-ECSA | 16.19 | 98.20 | 22997 | 1465.98 | 92.34 | 100 |
| SC5 | CHIKV | II-ECSA | 19.44 | 98.37 | 20486 | 1318.6 | 92.13 | 100 |
| SC6 | CHIKV | II-ECSA | 27.03 | 98.38 | 23933 | 1522.49 | 91.91 | 100 |
| SC7 | CHIKV | II-ECSA | 28.09 | 98.41 | 23517 | 1481.26 | 92.8 | 100 |
| SC8 | CHIKV | II-ECSA | 29.38 | 98.34 | 28991 | 1837 | 95.36 | 100 |
| SC9 | CHIKV | II-ECSA | 35.18 | 98.63 | 18274 | 1141.75 | 94.6 | 100 |
| SC10 | CHIKV | II-ECSA | 35.29 | 97.68 | 33821 | 2211.6 | 93.54 | 100 |
